# Proteomics and metabolomics reveal an abundant α-glucosidase drives sorghum fermentability for beer brewing

**DOI:** 10.1101/2022.06.30.498347

**Authors:** E. D. Kerr, G. P. Fox, B. L. Schulz

## Abstract

Sorghum (*Sorghum bicolor*), a grass native to Africa, is a popular alternative to barley for brewing beer. The importance of sorghum to beer brewing is increasing because it is a naturally gluten-free cereal and because climate change is expected to cause a reduction in the production of barley over the coming decades. However, there are challenges associated with the use of sorghum instead of barley in beer brewing. Here, we used proteomics and metabolomics to gain insights into the sorghum brewing process, to advise processes for efficient beer production from sorghum. We found that during malting, sorghum synthesises the amylases and proteases necessary for brewing. Proteomics revealed that mashing with sorghum malt required higher temperatures than barley malt for efficient protein solubilisation. Both α- and β-amylase were considerably less abundant in sorghum wort than in barley wort, correlating with lower maltose concentrations in sorghum wort. However, metabolomics revealed higher glucose concentrations in sorghum wort than in barley wort, consistent with the presence of an abundant α-glucosidase detected by proteomics in sorghum malt. Our results indicate that sorghum can be a viable grain for industrial fermented beverage production, but that its use requires careful process optimisation for efficient production of fermentable wort and high-quality beer.

## Introduction

Beer is one of the most popular beverages worldwide, with ∼1.95 billion hectolitres produced annually (1, 2). Typically, beer is made from malted barley seeds (*Hordeum vulgare* L. subsp. *vulgare*). While barley is central to current industrial brewing, sorghum (*Sorghum bicolor*) a grass native to Africa and the fifth most produced cereal worldwide (3), is a popular alternative to barley for brewing for several reasons. Depending on its eventual severity, climate change over the coming decades is expected to cause a reduction in the production of barley by 3–17% globally (4). This will have severe ramifications for beer production, price, and consumption. Sorghum is adapted to hot, drought-prone, semi-arid environments with low rainfall, making it well-suited for a warming climate. Global production of sorghum is therefore set to increase, making it an interesting alternative grain to barley for beer production (5). In addition, barley, wheat, rye, and oats are generally classified as gluten-containing cereals due to their prolamin storage proteins. Coeliac disease is caused by an inappropriate immune response triggered by the ingestion of gluten. ∼1% of the global population suffers from coeliac disease (6, 7), with this number increasing (8, 9). Many people are also gluten intolerant or choose to avoid gluten. Sorghum is inherently a gluten-free cereal, as its kafirin (prolamin) storage proteins do not trigger an immune response as in coeliac disease.

In many African countries sorghum is a staple food and is the main grain used to make fermented beverages (10). However, there are substantial shortfalls in the suitability of sorghum compared to barley for beer production. For the malting and brewing industries, the differences between sorghum and barley start at malting. Barley seeds are malted in three stages: steeping, germination, and kilning. Steeping is the process of repeatedly soaking the grains followed by periods of air-rest to drain the water, usually at 14-16 °C over 2-4 days (11), which increases grain moisture and stimulates metabolic activity. Excess water is removed and germination commences, allowing the synthesis of starch- and protein-degrading enzymes, usually over 3-6 days between 16-20 °C (11, 12). Seeds are then kilned to prevent excessive degradation of nutrients by the newly synthesised enzymes (1, 13–16). Malting of sorghum requires different conditions to barley, and malting parameters can impact sorghum malt quality. Amylases and proteases are lower in abundance in sorghum malt compared to barley malt (10, 17), but malting conditions can overcome these limitations. For example, in sorghum malting, lowering the degree of steeping (the percentage moisture content of the grain) to 41% compared to the standard ∼45% has been shown to increase free amino nitrogen (FAN) (18, 19). Barley is usually steeped for 24 – 48 h, while steeping sorghum for only 20 h improved α-amylase activity regardless of cultivar (10, 17, 20). Barley seeds are germinated anywhere from 3-7 days at 17-20 °C for malt production (20, 21). Sorghum germination of 4 days at 30 °C or 5 days at 26 °C results in the highest α- and β-amylase activity (17, 19). Germination temperatures of 20 °C and 25 °C have also been reported to result in similar amounts of reducing sugars and fermentable extracts (22). In summary, different malting conditions are needed for barley and sorghum, with the parameters for sorghum not yet standardised or optimised.

Malt is milled to open the grains and combined with warm water in the process of mashing to solubilise starch (gelatinisation) and proteins (solubilisation), and allow enzymes to degrade these macromolecules into fermentable sugars FAN (13, 15, 16, 23). The liquid fraction, wort, is separated from the spent grain and boiled with the addition of hops (*Humulus lupulus*). This hopped wort is cooled, fermented by yeast which utilise the soluble and consumable nutrients, matured, packaged, and sold. Mashing with sorghum requires 68–78°C for gelatinisation, significantly higher than 51–60°C used for barley (18, 24). The higher gelatinisation temperature of sorghum is likely due to its starch polymer structure (25, 26), and reduces the accessibility of enzymes to their starch substrate during mashing. Gelatinisation temperature is a key reason different mashing temperatures are used to mash sorghum malt and barley malt. The challenges of sorghum’s high gelatinisation temperature can be partially overcome by decantation style mashing, which involves boiling a part of the wort and adding it back to the main mash, resulting in higher extraction and fermentability than a standard infusion (single temperature) mash (22, 27). Brewing with sorghum requires optimisation of the malting and mashing process parameters, and even with optimised processes the wort produced will likely be less fermentable than barley wort.

Although sorghum proteins and starch have high solubilisation temperatures, its enzymes are comparatively heat sensitive (28). This unfortunate combination has a substantial impact on the quality of sorghum wort. Here, we used our established workflow for brewing proteomics (29) to gain insights into the potential deficiencies in the sorghum proteome compared to barley, to guide malting and mashing process design to increase the efficiency of wort production from sorghum (Fig. 1).

**Figure 1.**
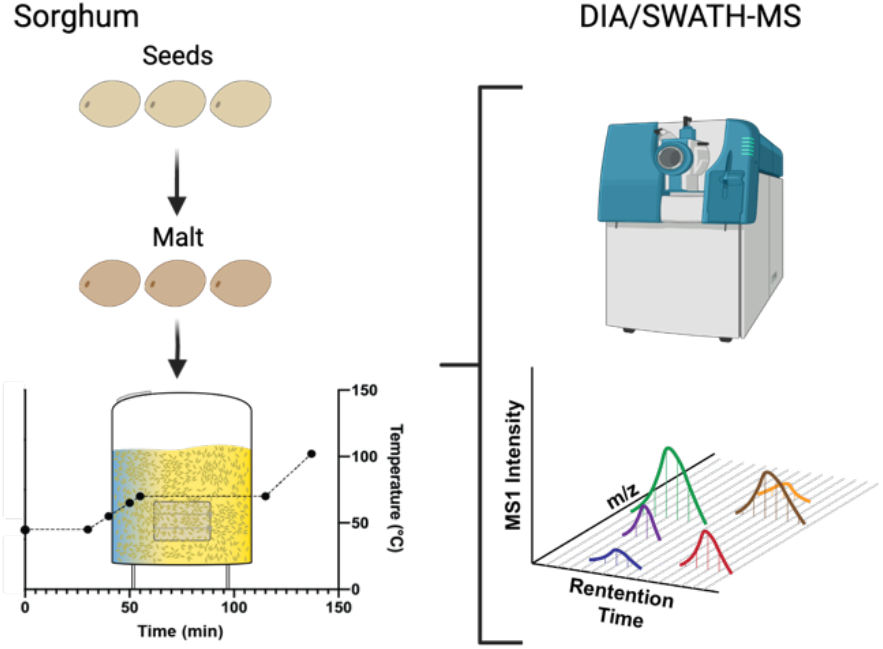
Overview of experimental design. Mature sorghum seeds were malted and mashes were performed for both sorghum malt and barley malt. Samples were collected of mature sorghum seeds, sorghum malt, and throughout both sorghum and barley mashes, and analysed by quantitative DIA/SWATH-MS proteomics.

## Methods

### Barley and sorghum malting

Micro-malting was carried out in a Phoenix automated micro-malting unit (Queensland Department of Agriculture, Leslie Research Facility, Toowoomba, Australia) to make fully modified malt as follows: 200 g barley grain (cultivar: Commander) was firstly steeped with water for 8 h, then 8 h air rest followed by another 8 h steeping, all at 17 °C. Following steeping, germination was allowed to occur by leaving at 17 °C for 4 days, turning barley grains every hour to avoid hot spots. 200 g sorghum grain (cultivar; Liberty) was steeped with water for 10 h, then 8 h air rest followed by another 8 h steeping, all at 21 °C. Following steeping, germination was allowed to occur by leaving at 21 °C for 4 days, turning sorghum grains every hour to avoid hot spots. For both barley and sorghum, germination was ceased by kilning: the grain was slowly heated to 50 °C to remove excess moisture, and then further heated to 80 °C for 24 h to suspend enzyme activity and to reduce the malt moisture to around 4%.

### Mash and boil experimental design

A small-scale mash was performed for both barley and sorghum malt in triplicate in Schott bottles incubating using an IEC programmable mash bath (IEC Melbourne, Australia). Mash method followed the Congress mashing program: 45 °C for 30 min and 70 °C for 60 min (30). Samples were taken at the start and end of both stages, and at 55 °C and 65 °C. Following mashing, wort was filtered and incubated at 102 °C for 5 min, after which a sample was taken.

### Proteomic sample preparation

Seed and malt samples were prepared as previously described (21, 31). Briefly, 10 mg of barley malt, 50 mg of sorghum seeds, or 50 mg of sorghum malt were ground and solubilized in 600 μL of 50 mM Tris-HCl buffer pH 8, 6 M guanidine hydrochloride, and 10 mM dithiothreitol (DTT) and incubated at 37 °C for 30 min with shaking. Cysteines were alkylated by addition of acrylamide to a final concentration of 30 mM and incubation at 37 °C for 1 h with shaking. Excess acrylamide was quenched by addition of an additional 10 mM DTT, samples were centrifuged at 18,000 rcf for 10 min, and proteins in 10 μL of the supernatants were precipitated by addition of 100 μL 1:1 methanol:acetone and incubation at − 20 °C overnight. Samples taken during the mash or boil were clarified by centrifugation at 18,000 rcf for 1 min and prepared as previously described (29). Briefly, proteins from 10 μL of barley wort or 30 μL of sorghum wort were precipitated by addition of 4 volumes 1:1 methanol/acetone and incubation overnight at − 20 °C. Precipitated proteins were resuspended in 100 μL of 100 mM ammonium acetate and 10 mM DTT with 0.5 μg trypsin (Proteomics grade, Sigma) and digested at 37 °C for 16 h with shaking.

### Proteomic mass spectrometry

Peptides were desalted with C18 ZipTips (Millipore) and measured by data dependent acquisition (DDA) and data independent acquisition (DIA) LC-ESI-MS/MS using a Prominence nanoLC system (Shimadzu) and TripleTof 5600 mass spectrometer with a Nanospray III interface (SCIEX) as previously described (32).

### Data analysis

Peptides and proteins were identified using ProteinPilot 5.0.1 (SCIEX). Sorghum samples were searched against all predicted proteins from *S. bicolor* v3.1.1 (PRJNA3869, downloaded 29 May 2017; 47,205 proteins) and barley samples were searched against all high confidence proteins from transcripts from the barley genome (GCA_901482405.1, downloaded 28 April 2017; 248,180 proteins). Both search databases also included contaminant proteins (custom database; 298 proteins). Search settings were: sample type, identification; cysteine alkylation, none; instrument, TripleTof 5600; species, none; ID focus, biological modifications; enzyme, trypsin; search effort, thorough ID.

The abundance of peptide fragments, peptides, and proteins was determined using PeakView 2.2 (SCIEX) with settings: shared peptides, allowed; peptide confidence threshold, 99%; false discovery rate, 1%; XIC extraction window, 6 min; XIC width, 75 ppm. Protein-centric analyses was performed as previously described (33), protein abundances were re-calculated using a strict 1% FDR cut-off of (21). Normalisation was performed to either the total protein abundance in each sample or to the abundance of trypsin self-digest peptides, as previously described (29). Principal component analysis (PCA) was performed using Python, the machine learning library Scikit-learn (0.19.1), and the data visualisation package Plotly (1.12.2). Protein and sample clustering was performed using Cluster 3.0 (34), implementing a hierarchical, uncentered correlation, and complete linkage. For statistical analysis, PeakView output was reformatted as previously described (21) and significant differences in protein abundance were determined using MSstats (2.4) (35) in R, with a significance threshold of p = 10^−5^ (36). Gene ontology (GO) term enrichment was performed using GOstats (2.39.1) (37) in R, with a significance threshold of p = 0.05 (36).

### Sugar and amino acid quantification by Multiple Reaction Monitoring

Post-boil samples were filtered, diluted 1:1000 in H2O, and measured by Multiple Reaction Monitoring (MRM) with UPLC-MS/MS as previously described (38). Sugars (glucose, maltose, and maltotriose) and amino acids (L-serine, L-proline, L-valine, L-threonine, L-leucine, L-isoleucine, L-aspartic acid, L-lysine, L-glutamic acid, L-methionine, L-histidine, L-phenylalanine, L-arginine, L-tyrosine, and L-cystine) were quantified using external calibration to multi-point standard curves with R^2^ values > 0.99.

### Fermentation assay

Fermentation of boiled wort was performed for both sorghum and barley wort using two brewing yeast strains: US-05 (Fermentis, American ale yeast) and M20 (Mangrove Jacks, Bavarian wheat yeast). Fermentations were tracked by weight loss as previously described (39) and samples were collected for ethanol quantification once ferments were complete.

### Ethanol quantification by headspace GC-MS/MS

Samples were diluted 1:50 in H2O, with 0.05% isopropanol added as an internal standard. Ethanol was quantified using external calibration to multi-point standard curves with R^2^ values >0.98. Chromatography was performed with a Rxi-624Sil 3.0 μm, 30 m x 0.53 mm column (Restek). Headspace GC-MS was performed as previously described (40), with a total run time of 20 min per sample.

### Data availability

The mass spectrometry proteomics data have been deposited to the ProteomeXchange Consortium via the PRIDE partner repository (41) with the dataset identifier PXD034981 (Username; Password: XiBUbGSV).

## Results and Discussion

### The proteome of malting sorghum mirrors malting barley

The malting process is essential to the brewing process, as it facilitates the synthesis of enzymes involved in degrading starch into fermentable sugars and protein into FAN. We aimed to investigate the biochemical changes that occur during malting of sorghum seeds, with a focus on the synthesis and abundance of key proteins important for the brewing process. To achieve this, we examined the proteomes of mature sorghum seeds and sorghum malt. DDA analyses identified 583 proteins, of which 533 were quantifiable by DIA/SWATH (Table S1 and S2). Principal component analysis (PCA) of the proteomic variance revealed that the proteomes of mature seeds and malt were clearly separated, indicating substantial proteomic changes during malting (Fig. 2A). We next performed statistical analysis to determine which proteins were significantly different in abundance between sorghum seeds and sorghum malt (Fig. 2B). We found 95 proteins that were significantly more abundant and 106 that were significantly less abundant in sorghum malt than in mature seed (Fig. 2B and Table S3).

**Figure 2.**
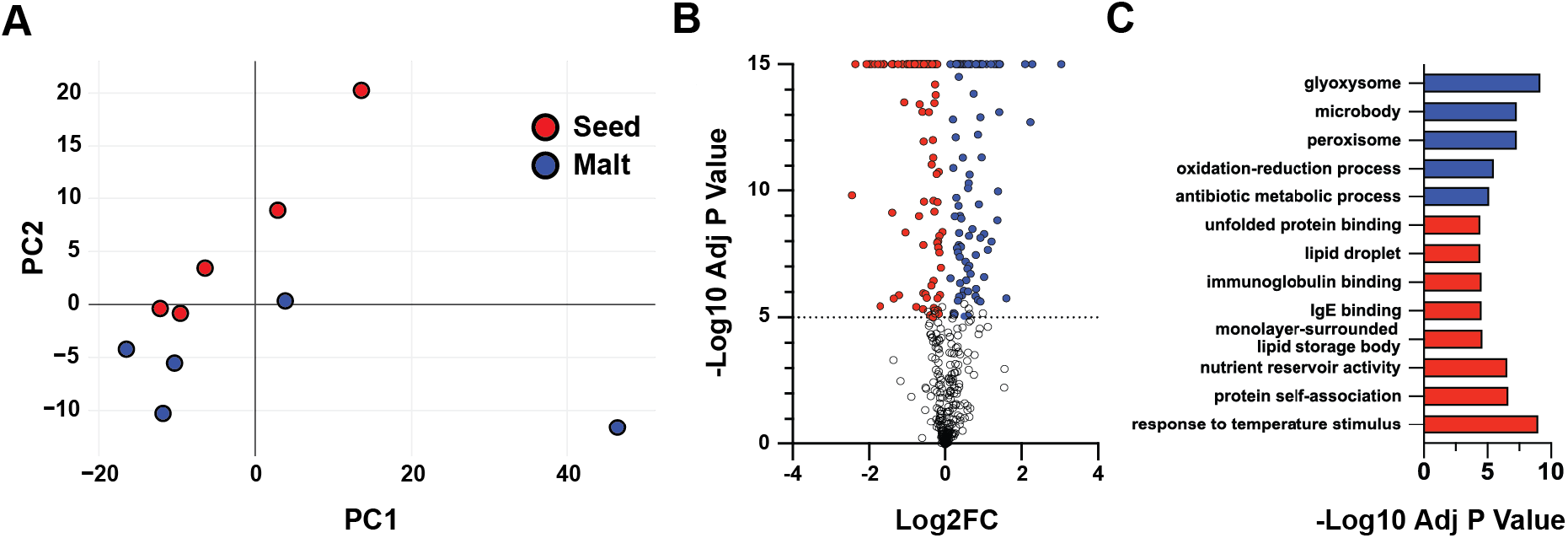
The impact of malting on the sorghum seed proteome. **(A)** PCA of protein abundance normalised to the abundance of trypsin in each sample. The first component (x-axis) accounted for 61.13% of the total variance and the second (y-axis) 15.30%. **(B)** Volcano plot of the comparison of mature sorghum seeds and sorghum malt. Grey, not significantly different; red, significantly (p < 10^− 5^) less abundant in malt; blue, significantly (p < 10^− 5^) more abundant in malt. **(C)** Significantly enriched GO terms for proteins with significant differences in abundance comparing mature seeds and malt. Values are shown as –log2 of Bonferroni corrected p-values for GO terms which were significantly enriched (p < 0.05) in proteins that were significantly more abundant in malt, blue; or significantly less abundant in malt, red.

Following statistical comparison between sorghum malt and mature sorghum seeds, we performed GO term enrichment analysis on proteins that were significantly different in abundance between malt and mature seeds (Fig. 2C and Table S4). A suite of GO terms were enriched in proteins significantly more abundant in sorghum malt than in mature sorghum seeds: “glyoxysome”, “microbody”, “peroxisome”, “oxidation-reduction process”, and “antibiotic metabolic process” (Fig. 2C). The enrichment of “glyoxysome”, “microbody”, and “peroxisome” was particularly interesting as they are all involved in the conversion of stored lipids into acetyl-CoA and then carbohydrates, which is common during germination (Fig. S1) (42, 43). As malt is partially germinated seed, this is consistent with an enrichment of “glyoxysome”, “microbody”, and “peroxisome” in the malt proteome.

A set of GO terms were enriched in proteins significantly less abundant in sorghum malt than in mature sorghum seeds: “response to temperature stimulus”, “protein self-association”, “nutrient reservoir activity”, “monolayer-surrounded lipid storage body”, “IgE binding”, “immunoglobulin binding”, lipid droplet”, and “unfolded protein binding” (Fig. 2C). Proteins contributing to “response to temperature stimulus”, “protein self-association”, and “unfolded protein binding” were heat shock proteins (HS177, HS181, HS232, HS23M, HS26P, HSP19, and HSP21) (Fig. S1), which have been previously shown to be abundant in mature barley seeds and which reduce in abundance after imbibition into germination in barley (21). Proteins which contributed to the GO terms “nutrient reservoir activity”, “IgE binding”, and “immunoglobulin binding” were seed storage proteins (11S2, CUCIN, GL19, GLB1, GLUB5, VCL21, VCL22, ZEG1) (Fig. S1), highlighting the abundance of seed storage proteins in mature sorghum seeds ready to be hydrolysed during germination. The reduction in abundance of seed storage proteins in sorghum malt aligns with the process of germination in barley, where seed storage proteins decrease in abundance during germination as they are digested by proteases resulting in smaller peptides and FAN (21). Proteins that contributed to “lipid droplet” and “monolayer-surrounded lipid storage body” GO terms were two oleosin proteins (OLE16 and OLEO3), an oil body-associated protein 1A (OBP1A), and a peroxygenase (PXG). Oleosins are known to help regulate oil body size and localisation (44), and oil body-associated proteins are known to be involved in stabilizing lipid bodies during desiccation (45), while peroxygenases are involved in the degradation of storage lipid in oil bodies during germination (46).

We found that some proteins were only able to be measured in either mature sorghum seeds or sorghum malt, but not both, indicating a dramatic change in protein abundance during malting. 22 proteins were unique to mature sorghum seeds, and 37 proteins were unique to sorghum malt (Fig. 3A). We performed GO term enrichment on these groups of uniquely measured proteins and found several GO terms enriched in proteins only present in sorghum malt (Fig. 3B). The most notable of these GO terms was “amylase activity”, which was contributed to by two α-amylases (AMY1 and AMY3C) and a β-amylase (AMYB). These amylases were not measured in mature seeds but were highly abundant in malt, consistent with their synthesis during the malting process (Fig. 3C). The appearance of amylases in malt is very promising for sorghum malting and mashing, as amylases are essential for producing fermentable sugars during mashing, which is needed for efficient downstream yeast fermentation.

**Figure 3.**
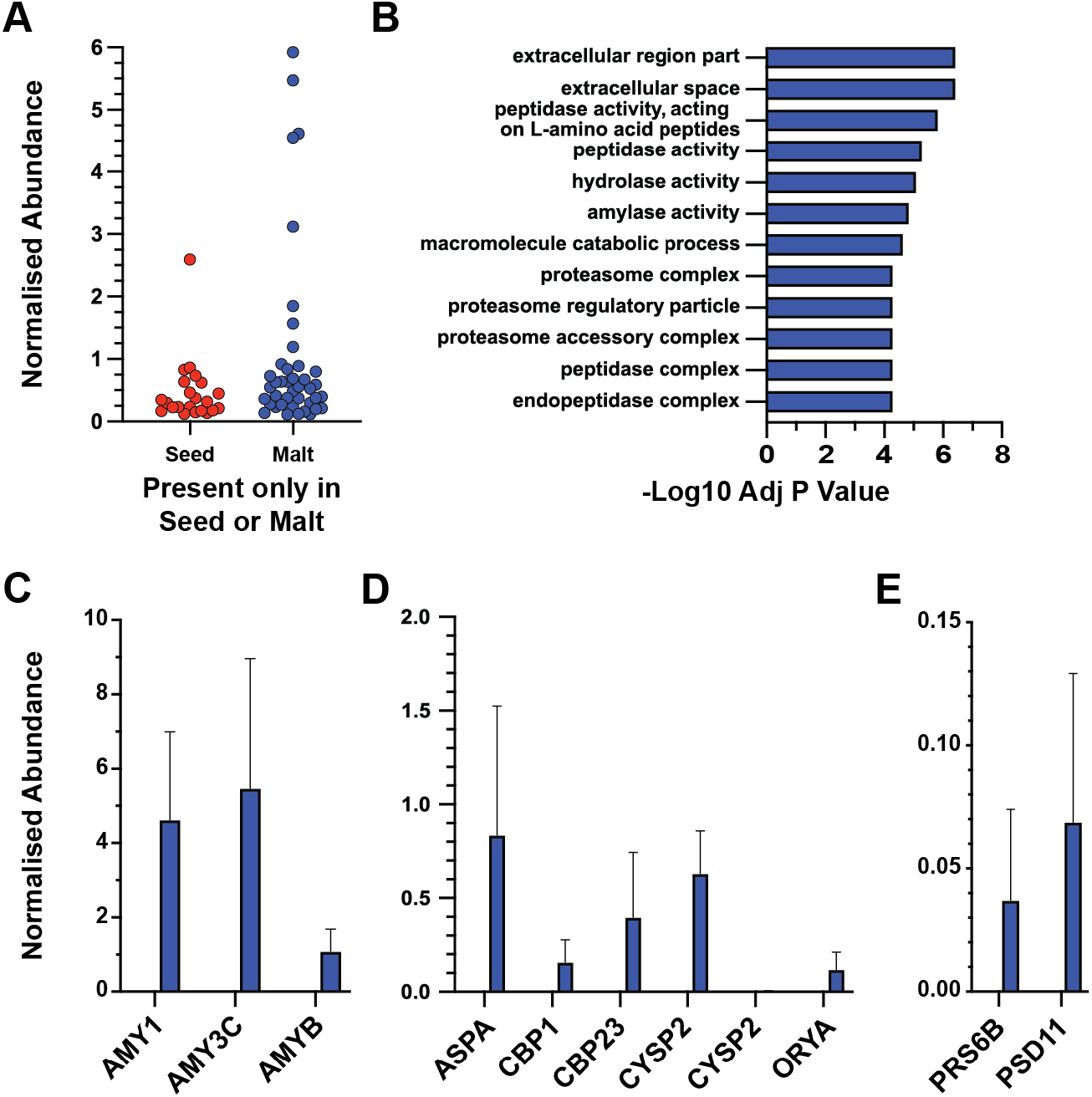
Sorghum germination/malting triggers synthesis of key brewing enzymes. **(A)** Abundance of proteins measured only in either malt or seed. Values, mean. Protein abundance normalised to the abundance of trypsin in each sample. **(B)** Significantly enriched GO terms in proteins unique to malt. Values are shown as -log_2_ of Bonferroni corrected p-values for GO terms which were significantly enriched (p < 0.05) in proteins that were only measured in malt. No GO terms were significantly enriched in proteins unique to seed. **(C)** Abundance of two α-amylase proteins; AMY1 and AMY3C and a β-amylase protein; AMYB. **(D)** Abundance of peptidases; aspartyl protease (ASPA), serine carboxypeptidase 1 (CBP1), serine carboxypeptidase II-3 (CBP23), two forms of cysteine proteinase EP-B 2 (CYSP2), and oryzain alpha chain (ORYA). **(E)** Abundance of proteasome proteins; 26S proteasome regulatory subunit 6B homolog (PRS6B) and 26S proteasome non-ATPase regulatory subunit 11 homolog (PSD11). Normalised protein abundance is relative to trypsin: values, mean; Error bars, SEM; Seed, red (no values shown). Malt, blue.

The enrichment of “peptidase activity, acting on L-amino acid peptides” and “peptidase activity” in proteins unique to malt was associated with proteases (ASPA, CBP1, CBP23, CYSP2, CYSP2, and ORYA) which were only found in sorghum malt (Fig. 3B and D). The increased abundance of proteases in sorghum malt aligned with the decrease in abundance of seed storage proteins, the main substrates of these proteases (13, 47). “Proteasome complex”, “proteasome regulatory particle”, and “proteasome accessory complex” were all associated with two proteasome regulatory subunits PRS6B and PSD11 (Fig. 3B and E). Proteasomes are protease complexes which assist in the degradation and recycling of specific proteins (48). Altered proteasome regulation may be associated with protein degradation to generate FAN or with regulation of the abundance of key proteins involved in germination. As with amylases, the synthesis of proteases during malting is promising for the performance of sorghum in malting and brewing, due to the requirement for yeast to be supplied with sufficient FAN for efficient fermentation.

The increase in abundance of amylases, proteases, and proteasome proteins along with the decrease in abundance of lipid droplet proteins, seed storage proteins, and heat shock proteins in sorghum malt compared to mature sorghum seeds is consistent with published proteomic changes in malted/germinating barley seeds (21). The conserved changes to the proteome of sorghum as it undergoes malting that we measured here indicate that sorghum malting shows all the hallmarks of standard industry barley germination (21).

### Nuanced differences in the dynamic mash proteome between sorghum and barley

Our proteomic analysis of mature sorghum seeds and sorghum malt showed that the nutrient proteins and enzymes needed for malting and mashing were present (Fig. 2 and 3). To better understand how sorghum malt performed in brewing, we performed side-by-side mash and boil of sorghum malt and barley malt, and investigated the proteome of the soluble wort fractions throughout the mash and boil (Table S1, S2, S5, and S6). Previously, we have shown that the barley mash proteome was dynamic and complex (29). We found that proteins increased in abundance early in the mash as they were extracted from the malt, then rapidly decreased in abundance at higher temperatures due to protein denaturation, aggregation, and precipitation due to lack of thermostability (29). Consistent with these processes, PCA of the wort proteome throughout the barley mash and boil showed a clear proteome shift as the mash progressed (Fig. 4A). In contrast, PCA of the wort proteome throughout the sorghum mash and boil showed a small shift in proteome through the early stages of mashing, with larger changes at higher temperatures (70 ºC – 60 min and 102 ºC) (Fig. 4B), likely because of the high protein solubilisation temperature in sorghum due to differences in starch polymer structure (18, 24).

**Figure 4.**
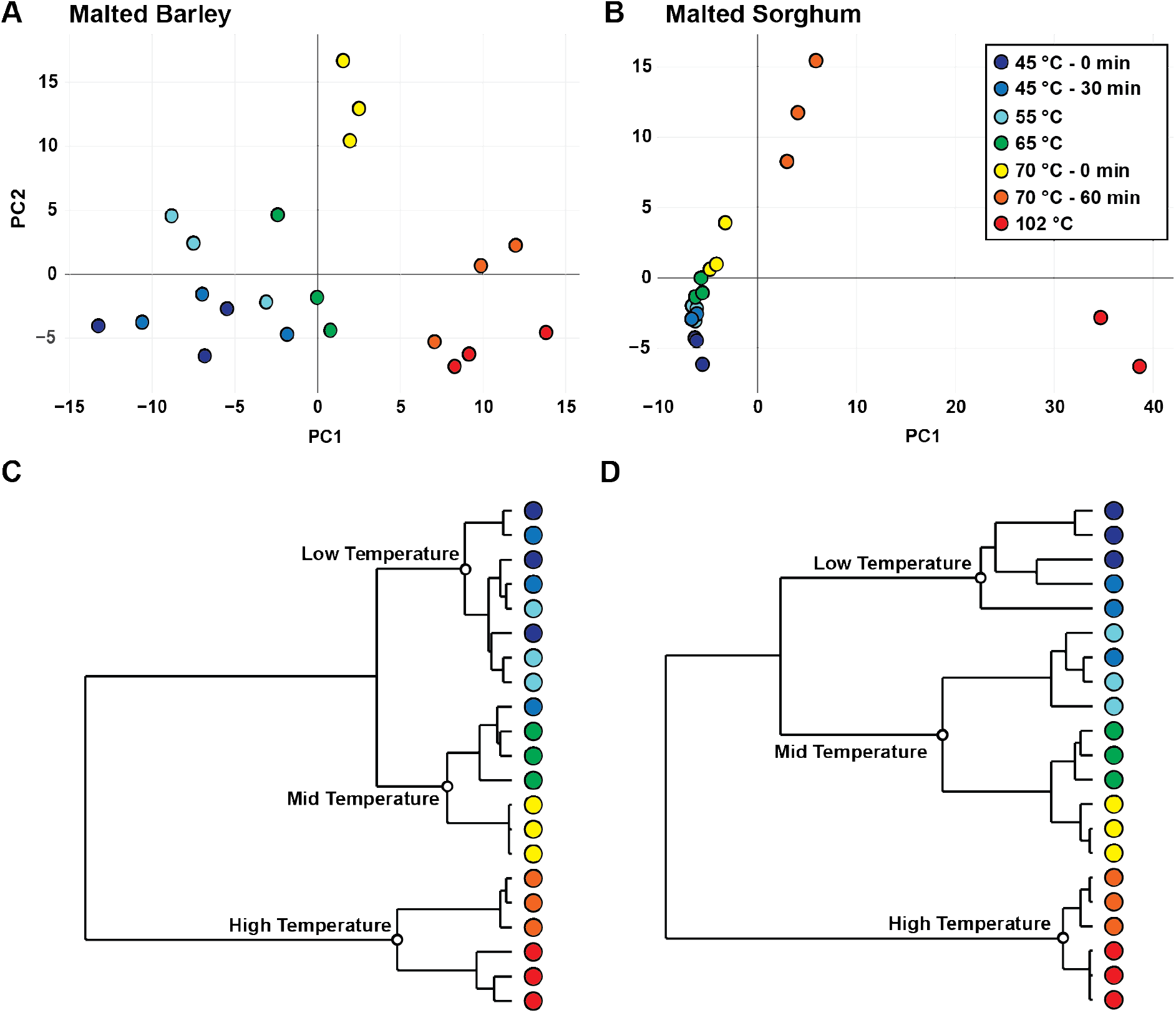
Delayed proteomic changes in wort in sorghum brewing compared to barley brewing. **(A)** PCA of barley mash and boil. The first component (x-axis) accounted for 36.36% of the total variance and the second (y-axis) 26.28%. **(B)** PCA of sorghum mash and boil. The first component (x-axis) accounted for 47.72% of the total variance and the second (y-axis) 9.13%. Dendrogram of **(C)** barley mash and boil and **(D)** sorghum mash and boil. Values were calculated from protein abundances normalised to the abundance of trypsin in each sample, clustered, and shown as both a PCA and a dendrogram. Clades visually noted within dendrograms were highlighted by white circles and labels.

To identify relationships between each stage of the barley and sorghum mash and boil, we next performed clustering analyses of the respective proteomes (Fig. 4C and D). In the barley mash, we noted three clades: low temperature (45 ºC and 55 ºC), mid temperature (65 ºC and 70 ºC – 0 min), and high temperature (70 ºC – 60 min and 102 ºC) (highlighted in Fig. 4C). Proteomes within the “low temperature” clade did not cluster by replicate, highlighting the limited overall proteomic changes occurring at low temperatures (Fig. 4C). The rapid change in proteomes at 65 ºC and above was highlighted by the separation of the “mid temperature” clade from the “low temperature” clade (Fig. 4C). The rapid shift in proteome from “low temperature” to “mid temperature” highlighted efficient protein extraction from barley malt as mash temperature increases. Finally, the separation of the “high temperature” clade from the other clades indicated a large shift in protein abundance as temperature increased and proteins began to denature, aggregate, and precipitate (Fig, 4C) (29). In the sorghum mash, as with barley, we identified three major clades, low temperature (45 ºC), mid temperature (55 ºC, 65 ºC, and 70 ºC – 0 min), and high temperature (70 ºC – 60 min and 102 ºC) (Fig. 4D). Within the “low temperature” and “mid temperatures” clades we saw a large overlap of replicates, suggesting a lack of substantial changes in the mash proteome during these stages (Fig. 4D). Towards the higher end of temperatures within the “mid temperature” clade and into the “high temperature” clade we saw stronger clustering of replicates. This clearer separation and clustering of “mid temperatures” indicated the start of protein extraction from sorghum malt, supported by the proteome variance in the PCA (Fig. 4B and D). The separation of the “high temperature” clade (Fig. 4D) that was also apparent by PCA (Fig. 4B) indicated a substantial shift in protein abundance as temperatures continued to increase and proteins potentially began to denature. Overall proteome dynamics throughout the mash and boil were similar for barley and sorghum malt, with the main apparent difference being that higher temperatures were required for protein extraction from sorghum malt.

The analysis of the overall variance in sorghum and barley brewing proteomes showed that while both showed qualitatively similar dynamics, the changes during the sorghum mash were muted in comparison to the substantial changes during the barley mash. The limited changes throughout the sorghum mash were potentially due to lower overall protein levels or to slower solubilisation from sorghum malt compared to barley. To better understand these differences, we compared the abundance of key proteins important to brewing performance: α- and β-amylase, seed storage proteins, and non-specific lipid transfer proteins (1, 13, 49). We found that in the barley mash, α-amylase behaved as expected, decreasing in abundance when temperature exceeded ∼65 ºC due to unfolding and aggregation (29) (Fig. 5A). The abundance of α-amylase was substantially different between mashes with the barley mash having an order of magnitude more α-amylase the sorghum mash (Fig. 5A and B). The abundance profile of α-amylase in the sorghum mash was also different to the barley mash, only increasing in abundance at 55 ºC and then decreasing at 70 ºC (Fig. 5B). β-amylase in the barley mash increased in abundance at a mash temperature of 55 ºC, followed by a substantial decrease in abundance at 65 ºC and eventual complete loss as the mash temperature rose further (Fig. 5C). Although we detected β-amylase in sorghum malt, it was not detected at any stage of the mash (Fig. 3C and 5D). The combined lower abundance of α- and β- amylase in the sorghum mash would likely affect overall brewing performance by severely limiting the production of fermentable sugars, causing reduced downstream fermentation efficiency and lower alcohol production (Fig. 5A – D).

**Figure 5.**
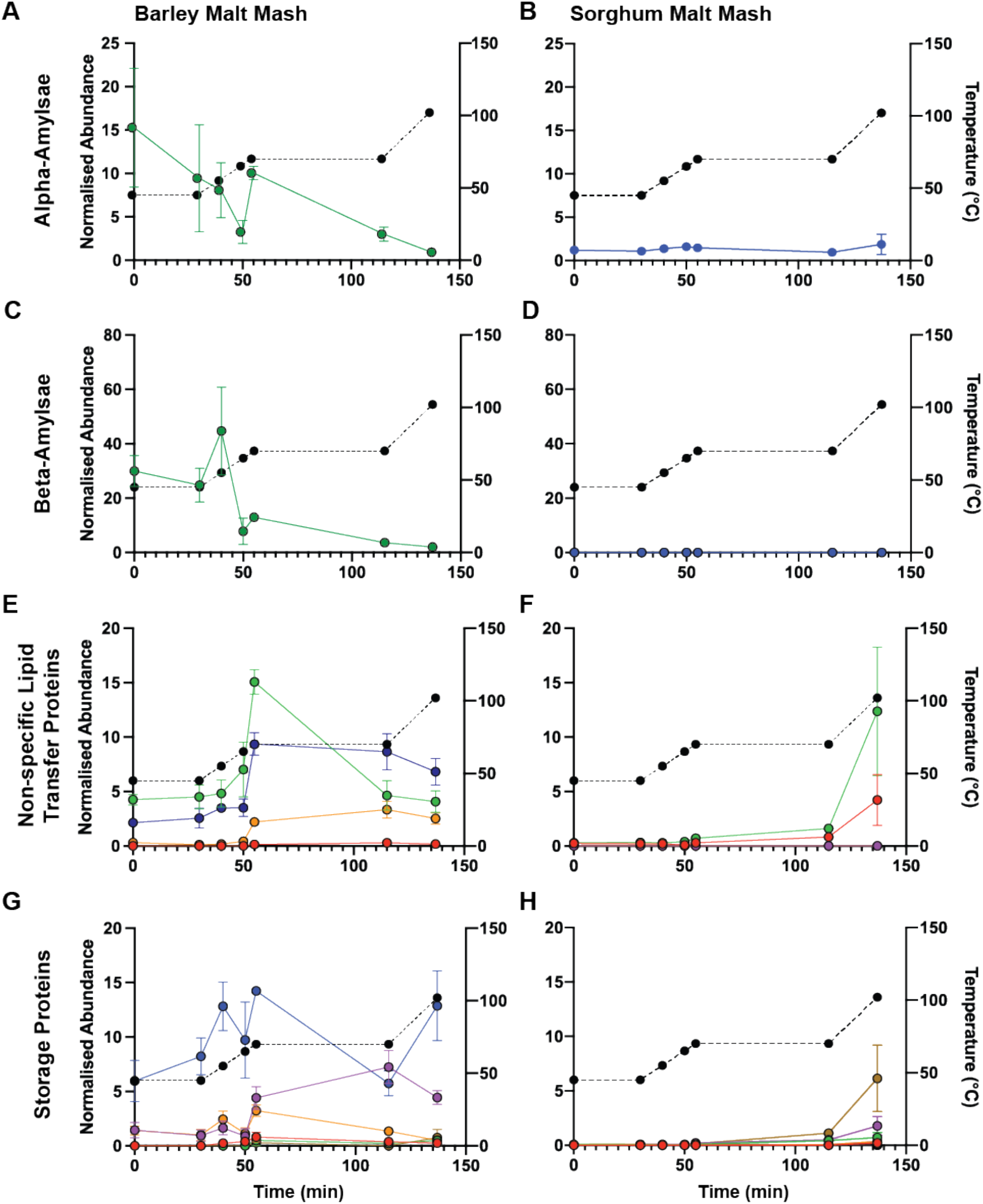
Dynamics of the abundance of key brewing proteins during the mash and boil for sorghum and barley. **A**bundance of summed forms of α-amylase in **(A)** barley and **(B)** sorghum. Abundance of summed forms of β-amylase in **(C)** barley and **(D)** sorghum. **(E)** Abundance of non-specific lipid transfer proteins in barley: NLTPX, red; NLTP2, green; NLTP1, purple; NLTP2, orange; NLTP4, blue. **(F)** Abundance of non-specific lipid transfer proteins in sorghum: NLT2G, red; NLTL1, green; NLTPX, light purple; NLTP2, orange; NLTP7, black; LTPG1, gold; NLTP2, blue; NLTP1, dark purple. **(G)** Abundance of Cupincin (CUCIN), red; Globulin-1 (GLB1), green; Vicilin-like seed storage protein At2g18540 (VCL21), purple; Vicilin-like seed storage protein At2g28490 (VCL22), orange; B3-hordein (HOR3), blue; Gamma-hordein-1 (HOG1), gold. **(H)** Abundance of 11S globulin seed storage protein 2 (11S2), red; Cupincin (CUCIN), green; 19 kDa globulin (GL19), light purple; Globulin-1 (GLB1), orange; Glutelin type-B 5 (GLUB5), black; Vicilin-like seed storage protein At2g18540 (VCL21), gold; Vicilin-like seed storage protein At2g28490 (VCL22), blue; 50 kDa gamma-zein (ZEG1), dark purple. A – H: Abundance (a.u; arbitrary units) of each protein normalised to trypsin. Values show mean, n=3. Error bars show SEM. Mash temperature profile is shown on right y-axis.

Lipid transfer proteins (LTPs) are important in beer as they are positively associated with foam formation and stability (50–52). In the barley mash, the abundance of LTPs increased at 55 ºC until the start of 70 ºC, and then decreased during the 70 ºC 60 min stage (Fig. 5E), consistent with previous reports (29). In the sorghum mash, the abundance of LTPs began to slowly increase only at 70 ºC and continued increasing throughout the remainder of the mash (Fig. 5F).

We next explored the dynamics of the abundance throughout the mash of seed storage proteins, which contain storage reserves of nitrogen, carbon, and sulphur for developing seeds (53). In brewing, FAN from seed storage proteins acts as nutrients for yeast instead of developing seeds, and mobilisation of these nutrients in the mash is critical for efficient yeast growth during fermentation (13, 54). In the barley mash, seed storage proteins increased in abundance at 55 ºC and then began to decrease in abundance at 70 ºC (Fig. 5E), as expected due to solubilisation and denaturation. In the sorghum mash, seed storage proteins began to increase slowly in abundance only at 70 ºC, reflecting either protein solubilization or starch gelatinisation only occurring at this high temperature.

The low abundance of seed storage proteins throughout most of the sorghum mash (Fig. 5H), in contrast to the high abundance of this class of protein in the barley mash (Fig. 5G) might limit their accessibility to digestion by proteases and result in low FAN production. Furthermore, barley seed storage proteins increased in abundance early in the mash but then decreased in abundance at higher temperatures (Fig. 5G) due to the relatively low thermal stability of proteolytically clipped forms (29). This decrease in abundance was not observed in the sorghum mash (Fig. 5H), also suggesting that protease activity was low throughout sorghum malting and mashing. To determine if we could detect direct evidence of protease activity in the sorghum mash, we inspected our peptide-level proteomic data for evidence of non-tryptic physiological proteolysis. We could indeed identify such physiological proteolytic events during the early stages of the mash (Fig. 6, and Tables S7 and S8). At lower mash temperatures in both barley and sorghum mashing, semi-tryptic peptides representing physiological proteolysis events were more abundant than full tryptic peptides representing un-proteolytically clipped protein (Fig. 6). As temperatures in the mash increased, the full tryptic peptides became the dominant form, reflecting the unfolding and aggregation of the proteolytically clipped protein forms (Fig. 6) (29). The abundance of semi-tryptic peptides in both sorghum and barley mashing suggested FAN was being created in either malting or the early stages of the mash in both cereals.

**Figure 6.**
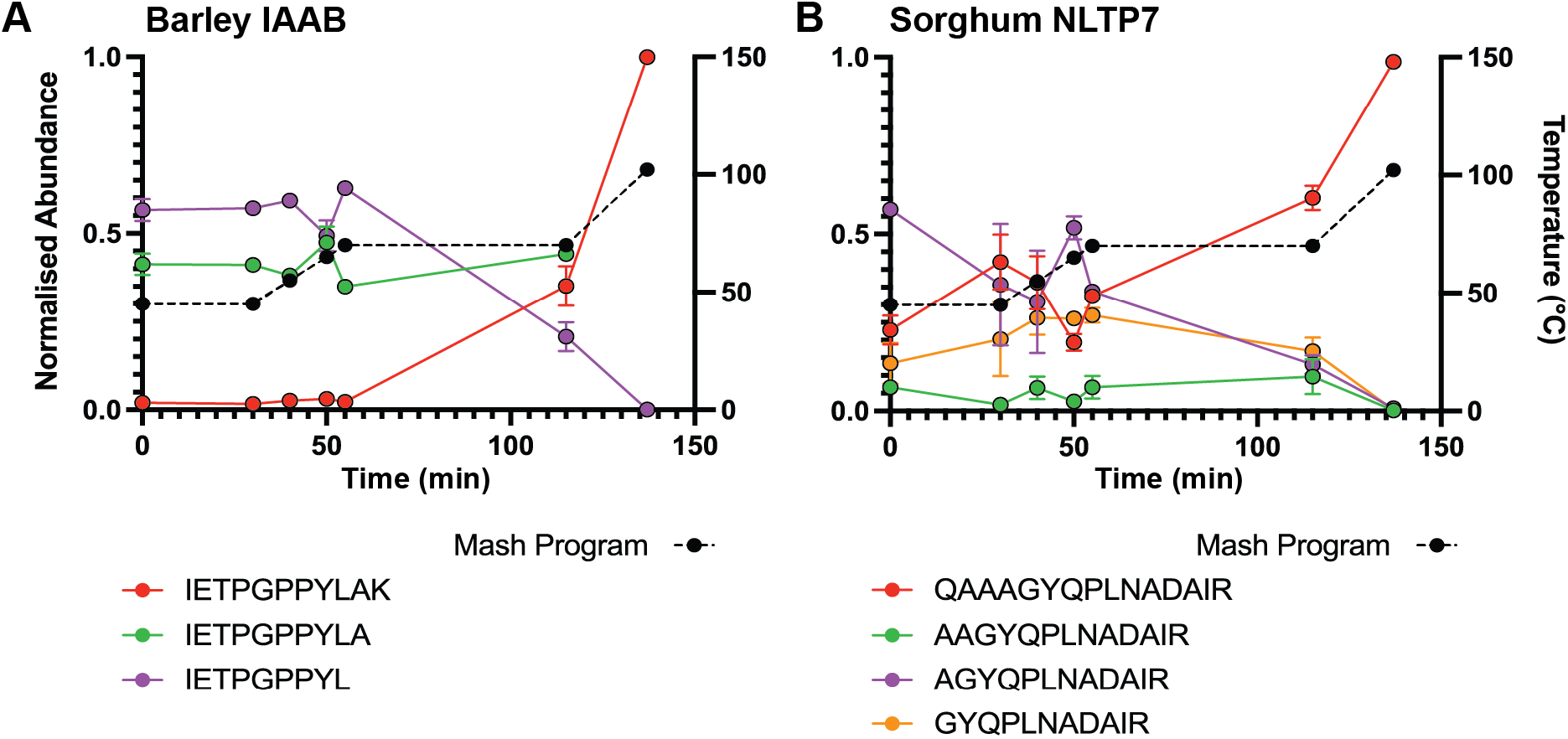
Abundant physiological proteolysis during the mash in both barley and sorghum. **(A)** Normalized abundance (a.u; arbitrary units) of full- and semi-tryptic peptides corresponding to R-I_55_ETPGPPYLAK_65_-Q from IAAB (α-amylase/trypsin inhibitor CMb) in barley **(B)** Normalized abundance (a.u; arbitrary units) of full- and semi-tryptic peptides corresponding to K-Q_87_AAAGYQPLNADAIR_101_-D from NLTP7 (Non-specific lipid-transfer protein 7) in sorghum. Values show mean, n=3. Error bars show SEM. Mash temperature profile is shown on the right y-axis.

### Comparable nutrient concentrations in sorghum and barley wort

Our proteomic analyses showed that while sorghum malt does contain the enzymes necessary for an efficient mash, they are present in low abundance and are only solubilised at much higher temperatures than from barley malt. We also found that wort produced with sorghum had a lower abundance of amylases and storage proteins, potentially due to sorghum malt having intrinsically lower levels of these proteins, or requiring a higher temperature for protein and starch solubilisation during mashing (Fig. 5). This may limit the fermentability of the final wort produced from sorghum. To determine the consequences of a lower abundance of amylases and other enzymes on sorghum wort quality, we used metabolomics to measure the amount of FAN and fermentable sugars present in barley and sorghum wort (Fig. 7A and B, and Table S9). Our proteomics data showed that in the sorghum mash, seed storage proteins that are critical sources of FAN were not highly abundant until late in the mash, likely compromising proteolysis and FAN production (Fig. 5F). However, we also found evidence of proteolysis in the early stages of mashing (Fig. 6B), consistent with the presence of at least some FAN-producing proteolytic activity. Using metabolomics, we measured the concentrations of 17 amino acids in sorghum and barley wort, and found significantly higher concentrations of L-alanine, L-lysine, L-histidine, and L-cystine in sorghum wort compared to barley wort, with no significant differences in the concentrations of the remaining 13 amino acids (Fig. 7A). This surprising result was consistent with our proteomic results, which showed proteolysis of seed storage proteins in sorghum malting or early in the mash, even though these proteins were not efficiently solubilized until very late in the mash. This is also consistent with previous studies of total FAN, which showed that regardless of protein abundance, both sorghum and barley wort have similar levels of FAN (55, 56).

**Figure 7.**
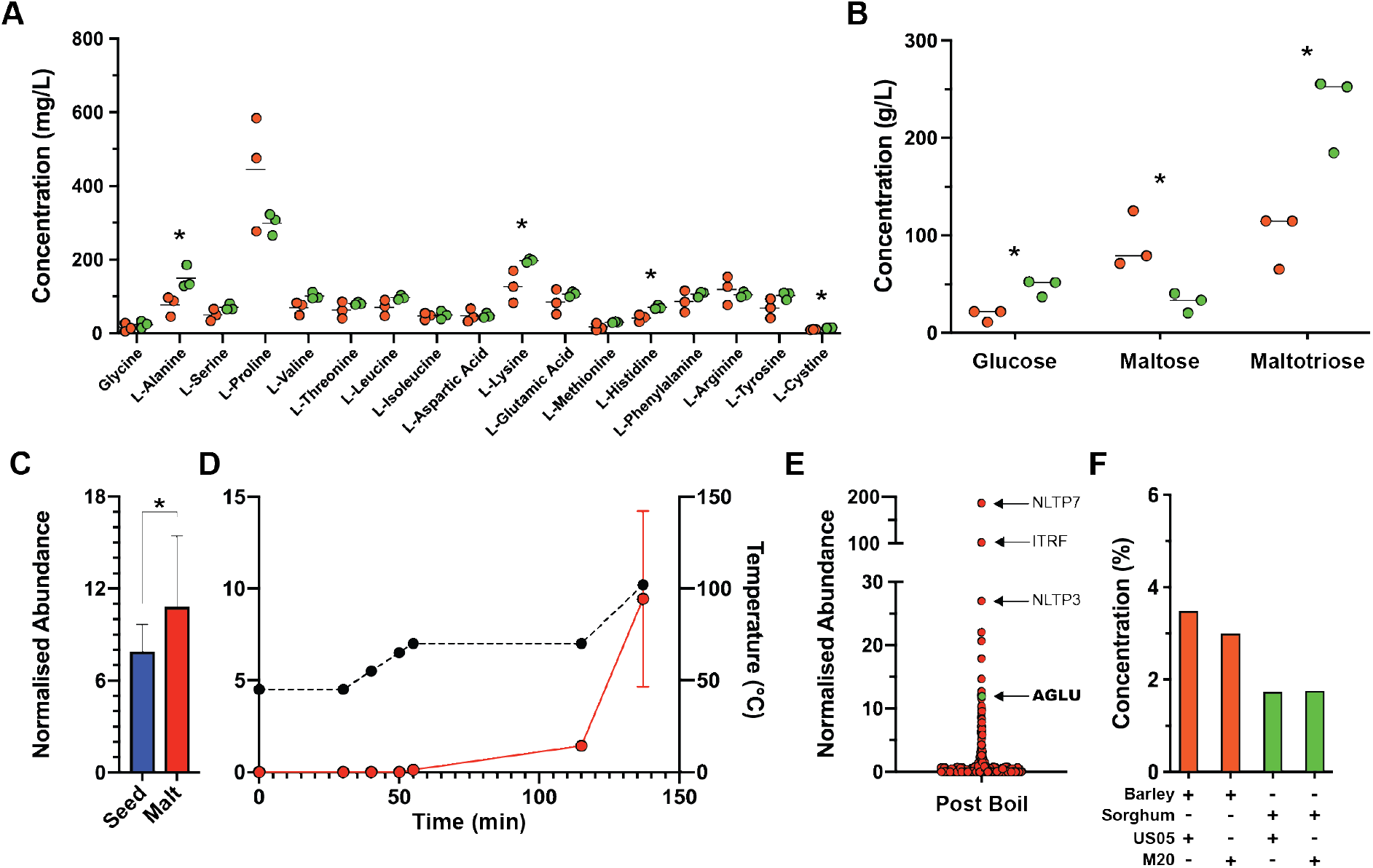
Fermentation performance of wort produced from barley malt and sorghum malt. **(A)** Amino acids quantified in wort produced from barley malt (orange) and sorghum malt (green). **(B)** Glucose, maltose, and maltotriose quantified in wort produced from barley malt (orange) and sorghum malt (green). Bars show mean, n=3. *, statistically significant differences between barley malt and sorghum malt (p < 0.05). Abundance of probable α-glucosidase (AGLU) in **(C)** sorghum seeds (blue) and malt (red), and **(D)** during the mash and boil. Values, mean; Error bars, SEM; *, p < 10^− 5^. **(E)** Abundance of proteins post boil. Values, mean. Proteins labelled: NLTP7 (Non-specific lipid-transfer protein 7), ITRF (Trypsin/factor XIIA inhibitor), NLTP3 (Non-specific lipid-transfer protein 3), and AGLU (probable alpha-glucosidase). **(F)** Concentration of ethanol produced in ferments from barley wort (orange) and sorghum wort (green), n=1.

We next compared the concentration of fermentable sugars present in wort from the sorghum and barley mashes. We found that there was significantly less maltose in sorghum wort (33.6 g/L) than in barley wort (79.1 g/L) (Fig. 7B). The low amount of maltose in sorghum wort was likely due to the limited abundance of α- and β-amylase in the mash (Fig. 6B and D). In contrast, the much higher level of maltose in barley wort was consistent with the abundant levels of α- and β-amylase throughout the mash (Fig. 6A and C). This difference in amylase abundance between barley and sorghum has been reported (28) and is likely a result of the inherent biology of barley as well as decades of breeding selecting for brewing performance. Despite the low abundance of α-amylase, we found that maltotriose concentrations in sorghum wort (252.5 g/L) were significantly higher than in barley wort (114.7 g/L) (Fig. 7B). Finally, and somewhat surprisingly, we found that there was significantly more glucose in sorghum wort (51.7 g/L) than in barley wort (21.8 g/L) (Fig. 7B). This relatively high glucose concentration is consistent with previous reports (55, 56). We therefore inspected our proteomic data to investigate if there was an enzyme present in the sorghum mash that could produce free glucose. We were able to identify a probable α-glucosidase (AGLU) that was not only abundant in sorghum malt but that was solubilised at later stages of the mash (Fig. 7C – E). In the final sorghum wort after the boil AGLU was the eleventh most confidently identified protein, consistent with high activity during mashing and potentially even during fermentation (Fig. 7E). This abundant α-glucosidase is likely the cause of significantly higher concentration of glucose and maltotriose in sorghum wort compared to barley wort (Fig. 7B).

While our proteomic analyses showed a low abundance of enzymes and seed storage proteins throughout the sorghum mash, our metabolomics analyses showed that sorghum wort had equivalent FAN and fermentable sugar profiles to barley wort. To functionally validate the suitability of sorghum and barley wort, we fermented the sorghum and barley wort with two commercial brewing strains of yeast (US05 and M20) and compared their fermentability (Fig. 7F and Table S10). We measured the extent of fermentation by tracking the weight loss caused by consumption of glucose, and production of CO2 and ethanol (39). Barley wort inoculated with US05 or M20 produced 3.48% or 2.99% ethanol (v/v), respectively (Fig. 7F). In contrast, fermentation of sorghum wort with US05 or M20 produced only 1.73% or 1.76% ethanol (v/v), respectively (Fig. 7F). Together with our mashing proteome data, this demonstrated that wort produced from sorghum malt using standard malting and mashing parameters was not as fermentable as wort produced from barley malt, but nonetheless resulted in efficient fermentation comparable to mid-strength beer.

## Conclusions and Future Directions

In this study we have shown that when it is malted, sorghum synthesises the enzymes needed for brewing, specifically amylases that degrade starch to fermentable sugars and proteases that digest seed storage proteins to produce FAN (21). However, our proteomic analyses of the dynamic sorghum mashing proteome highlighted that mashing with sorghum malt required higher temperatures for efficient protein solubilisation, and that the key enzymes α- and β-amylase were considerably less abundant in sorghum wort than in barley wort. This correlated with a lower amount of maltose in sorghum wort. Surprisingly, metabolomics analyses detected more glucose in sorghum wort than in barley wort, consistent with the presence of an abundant α-glucosidase in sorghum malt, AGLU. Finally, we showed that the fermentation of barley and sorghum wort was consistent with their FAN and fermentable sugar content, with efficient fermentation of barley wort and moderate fermentation of sorghum wort. Importantly, our results provide a molecular basis for previous descriptions of peculiarities and inefficiencies in sorghum wort production (10, 28).

Our results indicated that while sorghum mashing does not produce wort that is as fermentable as wort produced from barley mashing, it is still potentially viable for industrial beverage production. Our data showed that sorghum α-glucosidase AGLU is relatively stable at high temperatures and contributed to the high glucose concentrations in sorghum wort. The behaviour of this sorghum α-glucosidase and the high concentrations of maltotriose suggest that manipulation of sorghum mash parameters to allow α-glucosidase access to gelatinised starch for a longer period would produce wort that could support efficient fermentation.

## Supplementary Material

**Supplementary Figure 1.**
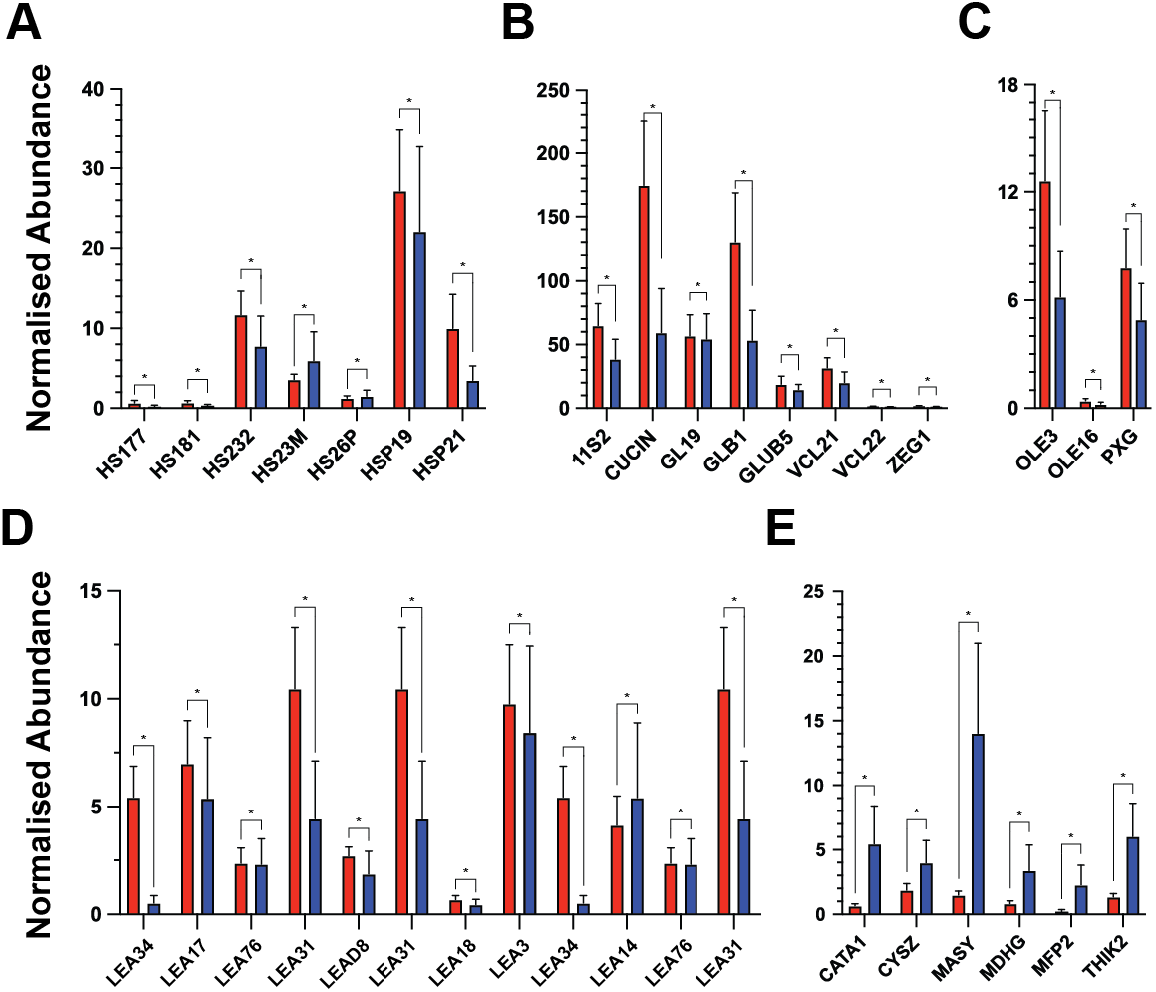
Abundance of germination related proteins in sorghum seeds and sorghum malt. **(A)** Abundance of heat shock proteins; HS177, HS181, HS232, HS23M, HS26P, HSP19, and HSP21. **(B)** Abundance of seed storage proteins; 11S2, CUCIN, GL19, GLB1, GLUB5, VCL21, VCL22, and ZEG1. **(C)** Abundance of lipid related proteins; OLE3, OLE16, and PXG. **(D)** Abundance of late embryogenesis proteins; two forms of LEA34, LEA17, two forms of LEA76, two forms of LEA31, LEAD8, LEA18, LEA3, and LEA14. **(E)** Abundance of glyoxysome related proteins; CATA1, CYSZ, MASY, MDHG, MFP2, and THIK2. For protein abundance: sorghum seeds, blue; sorghum malt, red. values, mean. Error bars, SEM. *, p < 10^− 5^.

